# Modeling binding of the conserved Csr/Rsm protein family across species of the γ-proteobacteria reveals niche-specific adaptation of the post-transcriptional regulon

**DOI:** 10.64898/2026.02.05.704111

**Authors:** Alexandra J Lukasiewicz, Lily G Hoefner, Ani Savk, Lydia M. Contreras

**Affiliations:** Department of Molecular Biosciences, The University of Texas at Austin, Austin, TX, United States; Department of Biology, The University of Texas at Austin, Austin, TX, United States; McKetta Department of Chemical Engineering, The University of Texas at Austin, Austin, TX, United States

**Keywords:** Post-transcriptional regulation, RNA-binding proteins, Computational biology, Ecology and evolution, Thermodynamic modeling

## Abstract

The γ-proteobacteria are an exceptionally diverse bacterial class whose members thrive in environments from deep-sea vents to human intestinal tracts. Rapid gene expression responses mediated by global post-transcriptional regulatory networks like the Csr/Rsm system are critical for bacterial survival in dynamic niches. CsrA/RsmA functions as a global regulatory RNA-binding protein, directly controlling hundreds to thousands of mRNA targets simultaneously across the transcriptome to coordinate systems-level metabolic and behavioral responses. Despite conservation of the CsrA/RsmA regulatory protein across γ-proteobacteria, the genes it regulates in different species remain poorly characterized. We extended a previously developed biophysical model of CsrA/RsmA-RNA binding from *Escherichia coli* and *Pseudomonas aeruginosa* to predict regulons across 16 diverse γ-proteobacterial species. While CsrA/RsmA protein structure and RNA-binding motif recognition are highly conserved, predicted target regulons diverge dramatically across species. Pathway enrichment analysis demonstrated both conserved regulation of core metabolic processes and extensive species-specific regulation of niche-adapted functions including virulence, biocontrol, and environmental stress response. Only two gene groups were shared exclusively among non-pathogens, while pathogens showed no exclusively conserved targets, indicating extensive regulon rewiring. These findings demonstrate that post-transcriptional regulatory networks evolve primarily through mutations in RNA targets that create or eliminate regulatory binding sites, rapidly adapting target repertoires to ecological demands while the regulatory protein mechanism remains conserved.

**Importance:** The CsrA/RsmA family represents one of the most influential global regulatory RNA-binding proteins in γ-proteobacteria, directly binding and regulating hundreds of mRNA targets to orchestrate systems-scale control over metabolism, virulence, and environmental adaptation, yet how this conserved mechanism adapts across diverse niches remains unclear. By predicting CsrA/RsmA targets across 16 species, we demonstrate that regulatory evolution occurs primarily through changes in targeted genes rather than the regulatory protein itself. This conserved mechanism with flexible targets may represent an efficient evolutionary strategy for optimizing gene expression for specific lifestyles, highlighting the importance of studying regulation beyond model organisms.

## 1.0 Introduction

Post-transcriptional regulation, particularly through direct RNA-binding protein (RBP) interactions with mRNA targets, is essential for bacterial adaptation to cellular and environmental stresses. Post-transcriptional regulators like CsrA/RsmA coordinate large-scale, systems-level regulatory effects by directly binding hundreds of mRNA targets. While CsrA/RsmA is conserved across gram-negative bacteria [1], regulating metabolism, motility, biofilm formation, and virulence [1, 2], the full spectrum of targets across species remains incompletely characterized. High-throughput techniques have advanced our knowledge [3–8], but many condition-specific or transient interactions remain undiscovered, and the evolutionary pressures shaping RBP-mRNA co-evolution are poorly understood, particularly in prokaryotes. The set of RNA targets that CsrA/RsmA directly regulates within an organism (which we refer to as its “regulon”) is thought to be shaped by the environmental niche [9, 10]. Based on this, we explore the hypothesis that specific environmental pressures and ecological niches drives diversification of global regulatory networks like Csr/Rsm.

To investigate the regulatory breadth of CsrA/RsmA across different bacterial species, we apply a previously developed biophysical model of Csr/Rsm binding [11, 12] to the transcriptomes of 16 members of the γ-proteobacteria that are representative of varying environmental niches (**Table 1**). These include pathogens of interest such as *Escherichia coli* O157:H7 EDL933, *Salmonella enterica* subsp. *Typhimurium, Legionella pneumophila, Legionella longbeachae, Pseudomonas syringae*, and *Vibrio cholerae*. We also include nonpathogenic organisms such as: *Escherichia coli* K-12 MG1655, *Azotobacter vinelandii, Aliivibrio fischeri, Halomonas hydrothermalis, Pseudomonas putida, Pseudomonas brassicacearum, Pseudomonas chlororaphis*, and *Pseudomonas protegens*. Much akin to DNA-binding transcription factors and eukaryotic RNA-binding proteins, CsrA/RsmA also exhibits high conservation across evolutionary distances while maintaining its systems-level regulatory capacity. Given the RNA binding rules that have been empirically derived for this protein [13, 14], its structural conservation [6, 13–17], and observed variation in regulatory response in different organisms [1, 4, 18], CsrA/RsmA represents a good model for interrogating evolutionary changes in regulatory RNA-protein interactions for prokaryotes. Our modeling approach offers a unique tool to explore the diversification of CsrA regulatory impact across species that allows for a deeper examination of these conserved regulatory protein-RNA interactions, revealing adaptations that may not be apparent from studying the evolutionary patterns of the CsrA/RsmA protein sequence alone. As part of this work, we first evaluate the suitability of our model towards capturing CsrA/RsmA-RNA interactions across species. We then discuss the overlapping and exclusive cellular pathways regulated across organisms and observe high variability in respective regulons, revealing that speciation and adaptation of the CsrA/RsmA post-transcriptional regulon is tailored to an individual niche.

**Table 1:** Overview of the 15 γ-proteobacterial species included in this study.

| Species | RsmA/CsrA Protein | % identity to E. coli CsrA | Major Pathways Regulated by CsrA/RsmA | Pathogenicity | Environment |
| --- | --- | --- | --- | --- | --- |
| *Halomonas hydrothermalis* | CsrA | 86.44 | Uncharacterized | non-pathogen | Deep-sea hydrothermal vents |
| *Vibrio cholerae* O1 biovar El Tor | CsrA | 94.74 | Iron metabolism, virulence, motility, lifestyle changes[7] | pathogen | Brackish saltwater, shellfish |
| *Aliivibrio fischeri* ES114 | CsrA | 90 | Symbiosis, siderophores, and luminescence [58] | non-pathogen | Marine environments, symbiosis with Eupryma scolopes |
| *Legionella longbeachae* NSW150 | CsrA | 76.67 | Uncharacterized | pathogen | Freshwater environments, hot tubs |
| *Legionella pneumophila* str. Paris | CsrA | 78.67 | Effector proteins, Type IV secretion system [59], intracellular growth [4] | pathogen | Intestinal tract, contaminated food and water |
| *Escherichia coli* O157:H7 str. EDL933 | CsrA | 100 | Pathogenesis, flagellar biosynthesis, cytochrome-c oxidases, nitrogen metabolism[33] | pathogen | Contaminated food and water |
| *Escherichia coli* K-12 MG1655 | CsrA | 100 | Motility, glycogen synthesis, biofilm formation[1] | non-pathogen | Laboratory strain |
| *Salmonella enterica* serovar Typhimurium | CsrA | 100 | Virulence, metabolism, and biofilms[1] | pathogen | Intestinal tract, contaminated food and water |
| *Azotobacter vinelandii* DJ | RsmA | 86.69 | Alginate and polymer biosynthesis [60] | non-pathogen | Agricultural soils |
| *Pseudomonas aeruginosa* UCBPP PA14 | RsmA | 82.65 | Lifestyle changes, virulence, secretion systems, quorum sensing, biofilms [1] | pathogen | Ubiquitous, soil, water, clinical settings |
| *Pseudomonas brassicacearum* subsp. *brassicacearum* | RsmA | 75.81 | Secondary Metabolism, Plant hormone synthesis, secretion systems, biofilms [61] | non-pathogen | Rhizosphere of brassica crops |
| *Pseudomonas syringae* pv. *Tomato str DC3000* | RsmA | 75.81 | Secretion systems, motility, pathogenesis, siderophores[36] | pathogen | Plant leaf surfaces |
| *Pseudomonas fluorescens* 1206 | RsmA | 75.81 | Secondary metabolism, secretion systems, antifungal compound expression [62] | non-pathogen | Agricultural soils, rhizosphere |
| *Pseudomonas putida* KT2440 | RsmA | 75.41 | Motility, biofilm formation, stress response [63, 64] | non-pathogen | Agricultural soils |
| *Pseudomonas chlororaphis* subsp. *aureofaciens* 30-84 | RsmA | 75.47 | Phenazine production, quorum sensing, biofilm formation[65] | non-pathogen | Wheat rhizosphere |
| *Pseudomonas protegens* Pf-5 | RsmA | 75.47 | Secondary metabolites, motility and oxidative stress [25] | non-pathogen | Plant rhizosphere |

## 2.0 Results

### 2.1 Evaluation of motif energetics across modeled species reveals conservation of binding motif

A two-state thermodynamic model of CsrA protein binding was previously constructed by our group to model direct interactions with mRNAs in *E. coli* [11] and the homologous RsmA protein interactions across the *P. aeruginosa* transcriptome [12]. Given the transferability of the model between these two species, and the complementation of CsrA activity in other organisms [19, 20], we hypothesized that our initial model could be applied to other members of the γ-proteobacteria.

Alphafold3 modeling [21] was used to generate structural predictions of the native CsrA protein from 10 species: *P. aeruginosa, P. protegens, A. vinelandii, L. pneumophila, L. longbeachae, V. cholerae, A. fischeri, S. enterica, E. coli*, and *H. hydrothermalis* in complex with the *hcnA* stem-loop RNA sequence. To evaluate structural similarity to the experimentally-derived crystal structure of the *P. protegens* RsmE-*hcnA* complex (PDB: 2JPP), the Template Modeling (TM) score were calculated for the predicted structures using the TM-score tool [22]. TM scores were all greater than 0.5, indicating a high similarity. Alphafold3-generated PDB structures were then used to generate species-specific positional weight matrices (PWMs) that measure the energetic contribution of each individual nucleotide in a given motif to binding affinity. We then generated each PWM given the relative change in binding affinity using the Rosetta-Vienna RNP ΔΔG tool [23] as described in [12]. ΔΔG values correlated with the RsmE-*hcnA* PWM [12] for all organisms (**Figure 1**). The calculated energy scores and representative motifs can be found in **Supplementary Table 3**. Given the high correlation between energies for each species and the conservation of protein structure, we concluded that the model generated for evaluating RsmA binding in *P. aeruginosa* described in [12] was applicable across multiple genomes.

**Figure 1:**
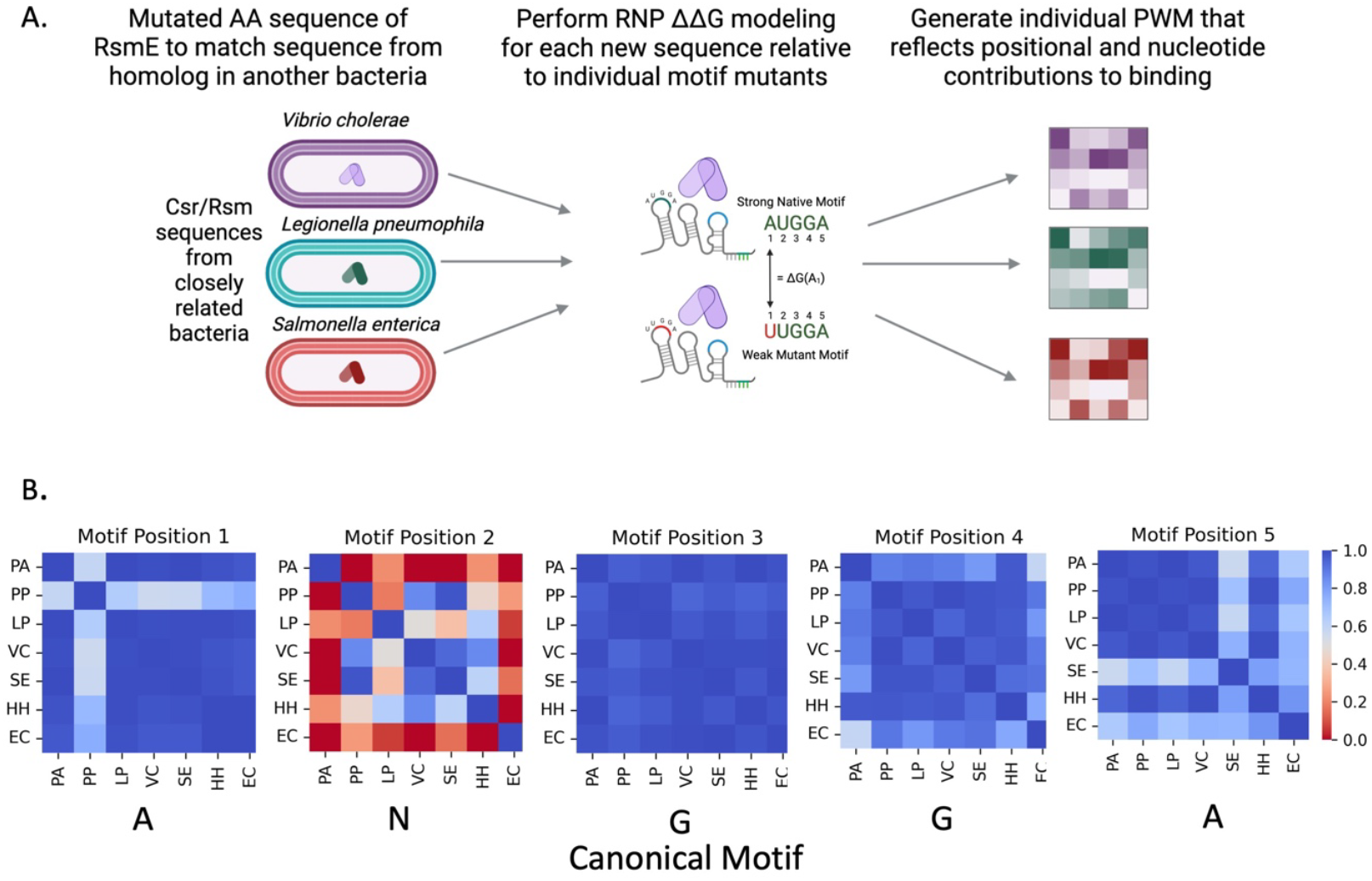
Process of PWM generation and evaluation for species modeled. A) Overview of RNP ΔΔG modeling procedure used to generate PWMs per organism. B) All vs. all correlation plots of the predicted ΔG-nt at each position reveals core ANGG conservation, with some variability observed for members of the Enterobacteriaceae in position 5. PA- *P. aeruginosa*, PP- *P. protegens*, LP- *L. pneumophila*, VC- *V. cholerae*, SE- *S. enterica*, HH- *H. hydrothermalis*, EC – *E. coli*

### 2.2 Model predictions and energetic cutoffs are validated against experimental data

To validate our biophysical thermodynamic model and establish effective filtering cutoffs for distinguishing direct binding targets (genes with CsrA/RsmA binding sites leading to regulatory effects) from indirect targets (genes regulated downstream through regulatory cascades or with weaker, non-functional binding interactions), we compared predictions against published high-throughput datasets for species with experimental CsrA/RsmA characterization: *E. coli* K-12 MG1655 and O157:H7 EDL933 [18, 24], *S. Typhimurium* [8], *L. pneumophila* [4], *V. cholerae* [7], and *P. protegens* [25]. Prior computational screens by [26] predicted direct targets in several of these species using sequence motif finding, and were also used to validate our model. Unlike binary sequence motif searches, our model generates continuous affinity scores (ΔG_total_) for each gene transcriptome-wide. We employed hypergeometric enrichment testing to identify species-specific energetic cutoffs that optimally discriminate experimentally validated targets from non-targets, following the approach validated for *P. aeruginosa* RsmA [12]. Since median ΔG_total_ distributions differed significantly between species (Kruskal-Wallis p < 2.2×10^-16), predicted energies were scaled 0-1 to enable cross-species comparison and establish a generalizable cutoff for uncharacterized organisms.

Hypergeometric enrichment testing established species-specific cutoffs: *E. coli K-12* (0.73, 750 targets/21.5% of transcriptome), *E. coli EHEC* (0.69), *S. enterica* (0.67, 431 targets/10%), *L. pneumophila* (0.56, 1,446 targets/46.8%), *V. cholerae* (0.50, 835 targets/23.7%), and *P. protegens* (0.61, 1,130 targets). Averaging species-specific cutoffs from all validated organisms we derived a cross-species cutoff of 0.62 ± 0.098 for these and other less-characterized species (**Figure 2**). Applying this cutoff, the majority of known biochemically validated targets were correctly predicted in all species (**Figure 2**) with some exceptions, indicating our cross-species cutoff is highly conservative.

**Figure 2.**
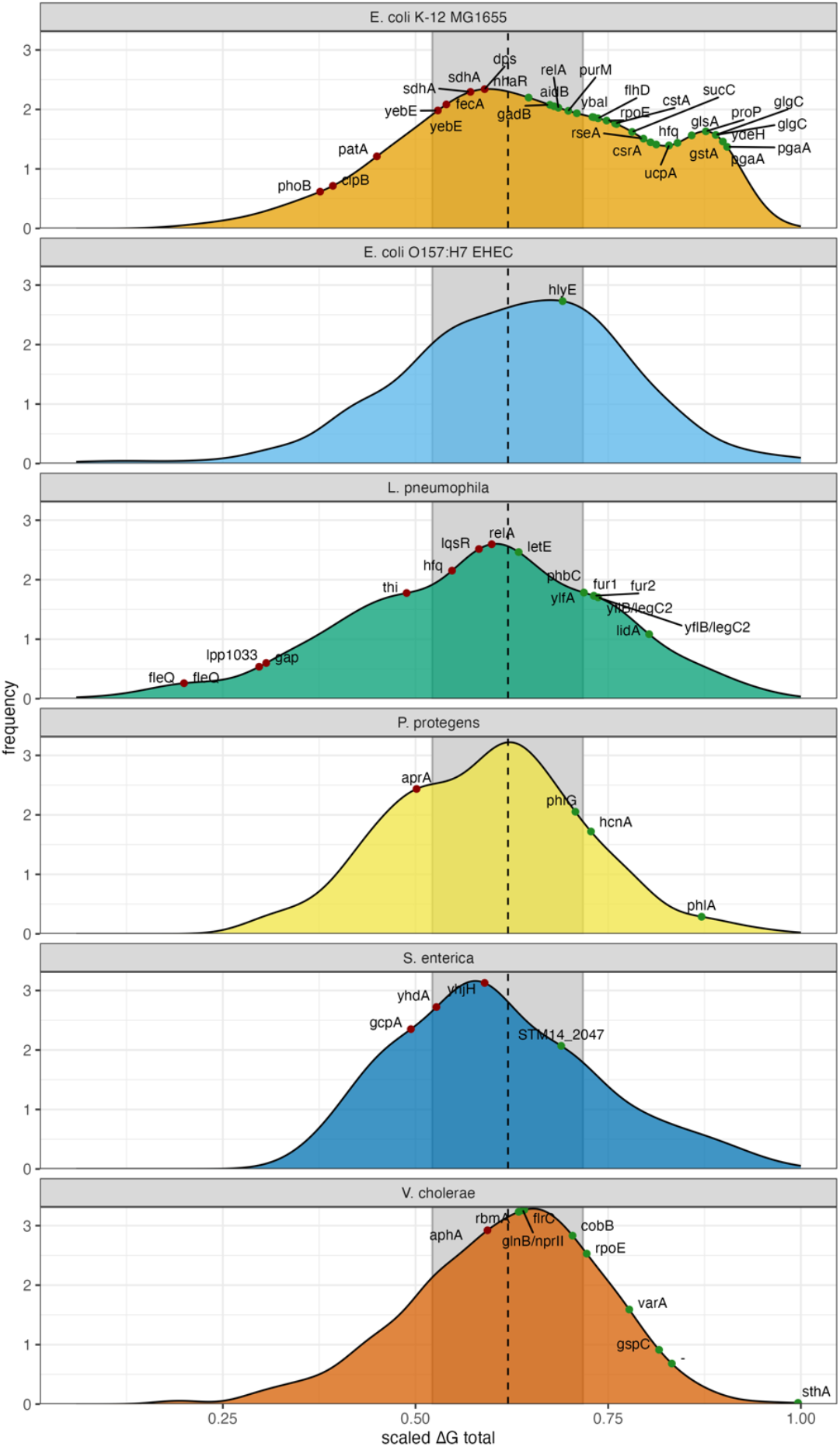
Distribution of a scaled total affinity scores (ΔGtotal) for species with prior experimental characterization of CsrA binding validates predictive capabilities of the model. Dashed black line indicates averaged universal cutoff of 0.62 to differentiate a target (green) from a non-target (red). In *E. coli K-12 MG1655*, known direct targets including *glgC, pgaA*, and *flhD* were correctly identified, while negative controls (*phoB, fecA* [66]) were appropriately excluded. Several genes with prior evidence of direct binding (*clpB, patA, yebE, sdhA, dps*) did not pass the cutoff, indicating conservative filtering that may yield false negatives. For *E. coli* O157:H7 EDL933 the biochemically validated *hlyE* leader [24] passed our universal cutoff value. In *S. enterica*, characterized targets of CsrA were correctly predicted, however our general cutoff value excluded *gcpA, yhdA*, and *yhjH* (Figure 3). All known direct targets in *V. cholerae* except *aphA* were captured; *aphA* would be included with a *V. cholerae*-specific cutoff but fell below our conservative generalized threshold.

These validations demonstrate that our model accurately captures direct CsrA/RsmA binding interactions across phylogenetically diverse γ-proteobacteria, with species-specific cutoffs accommodating biological variation in regulatory breadth. The conservative nature of our cutoffs (evidenced by missed targets like *clpB, patA, aphA*) prioritizes specificity, providing high confidence in subsequent comparative analyses while acknowledging some false negatives. All predicted targets are summarized in **Supplementary Table 1**.

### 2.3 Lack of shared core regulon is present across modeled species

To systematically compare CsrA/RsmA regulation across evolutionary distances we mapped protein coding genes to orthogroups using the EggNOG-mapper v2 tool [27]. Orthogroups represent clusters of genes descended from a single ancestral gene, allowing us to track regulatory conservation across evolutionary distances while accounting for gene duplications and losses.

Enrichment testing was performed to assess whether predicted CsrA-targeting orthogroups are conserved or divergent across these modeled species. We hypothesize that significant overlap of predicted targets would therefore indicate a core regulon that does not diverge depending upon the organism. Hierarchical clustering of overlapping orthogroups reveals that major enrichment is found at the family level (**Figure 3A**), indicating that the most significant enrichment of overlapping targets is shared within each family taxonomic designation. The degree of significant enrichment was not shared across all modeled species (**Figure 3A**), indicating that CsrA regulation may be tuned to address the specific metabolic needs, stresses, and environmental niches that each species inhabits.

**Figure 3:**
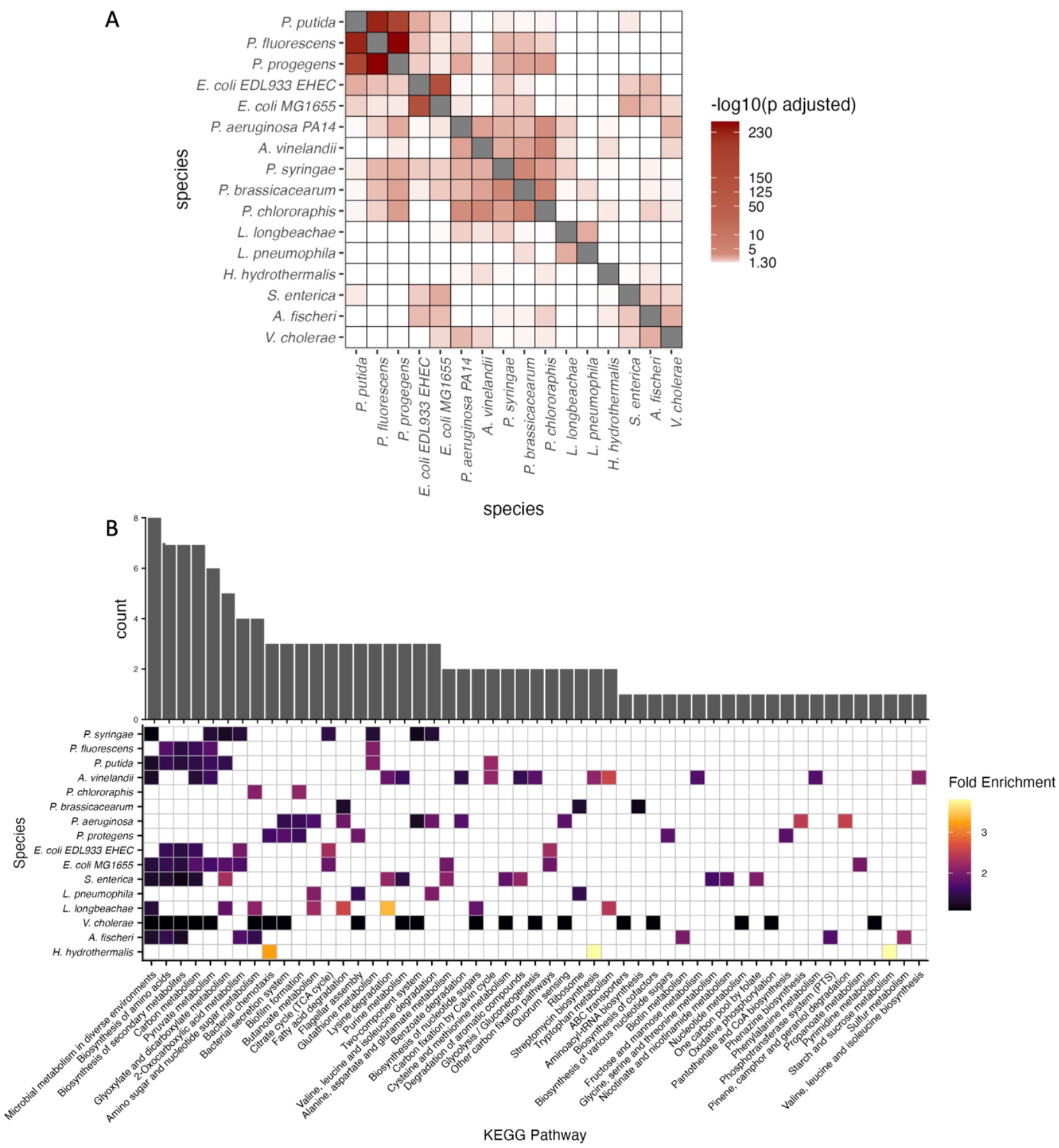
Predicted CsrA regulation of eggnog mapped orthogroups reveals small handful of conserved targets. A) Hypergeometric overlap testing shows significant overlap of CsrA targets occurs on the family level, but is not shared broadly across all species in the γ-proteobacteria. B) Overlapping and exclusive KEGG pathways represented per species following enrichment analysis. Some core pathways appear shared on the genus level, but each species exhibits some unique distribution of processes regulated. Genes involved in each pathway can be found in **Supplementary Table 1**

Despite CsrA/RsmA functioning as a global regulator of gene expression, we observed remarkably limited conservation of specific targets across species. No single orthogroup contained shared CsrA/RsmA targeting across all 16 modeled species, indicating the absence of a core regulon. Three orthogroups, aldehyde dehydrogenases (1RMBQ), cold-shock proteins (1SCA7), and phage integrases (1RMJ1) were predicted to be targeted within 15 of the 16 bacteria (**Supplementary Table 1**).

We next examined whether pathogenicity correlates with shared regulatory targets. Suprisingly, pathogenic species share no exclusive targets of CsrA/RsmA, while nonpathogens share targeting of two unique orthogroups: the group encoding for dTDP-glucose 4,6-dehydratase, *rfbB* (1RP7G) (**Supplementary Figure 1**), and the gene encoding for small protein B *smpB* (1S3PT) (**Supplementary Figure 2**). This asymmetry suggests that pathogenic lifestyles may require more diverse, species-specific regulatory adaptations.

### 2.4 CsrA/RsmA maintains core metabolic regulation while enabling niche-specific functional adaptation

To characterize the functional impact of CsrA regulation across diverse γ-proteobacteria, we performed enrichment analyses of predicted CsrA target-orthogroups to KEGG pathways, Gene Ontology (GO) terms, Clusters of Orthologous Groups (COG), and Pfam domains. These analyses revealed both conserved and niche-specific regulatory patterns, highlighting the dual role of CsrA in maintaining core cellular functions and enabling environmental adaptation.

KEGG pathway enrichment analysis across all 16 species revealed both deeply conserved regulation of central metabolism (**Figure 3B**). Core metabolic pathways including carbon metabolism, glycolysis/gluconeogenesis, pyruvate metabolism, and the TCA cycle were enriched across nearly all species, indicating that CsrA/RsmA consistently regulates metabolic flux regardless of ecological niche.

COG enrichment analysis confirmed this pattern, showing that nucleotide transport and metabolism (COG J, F) were commonly regulated across species (**Figure 4A–B**). However, the regulatory scope extends far beyond these core functions, with extensive species-specific diversification reflecting adaptation to distinct lifestyles..For example, *L. longbeachae* uniquely showed enrichment in defense mechanisms (COG V).

**Figure 4:**
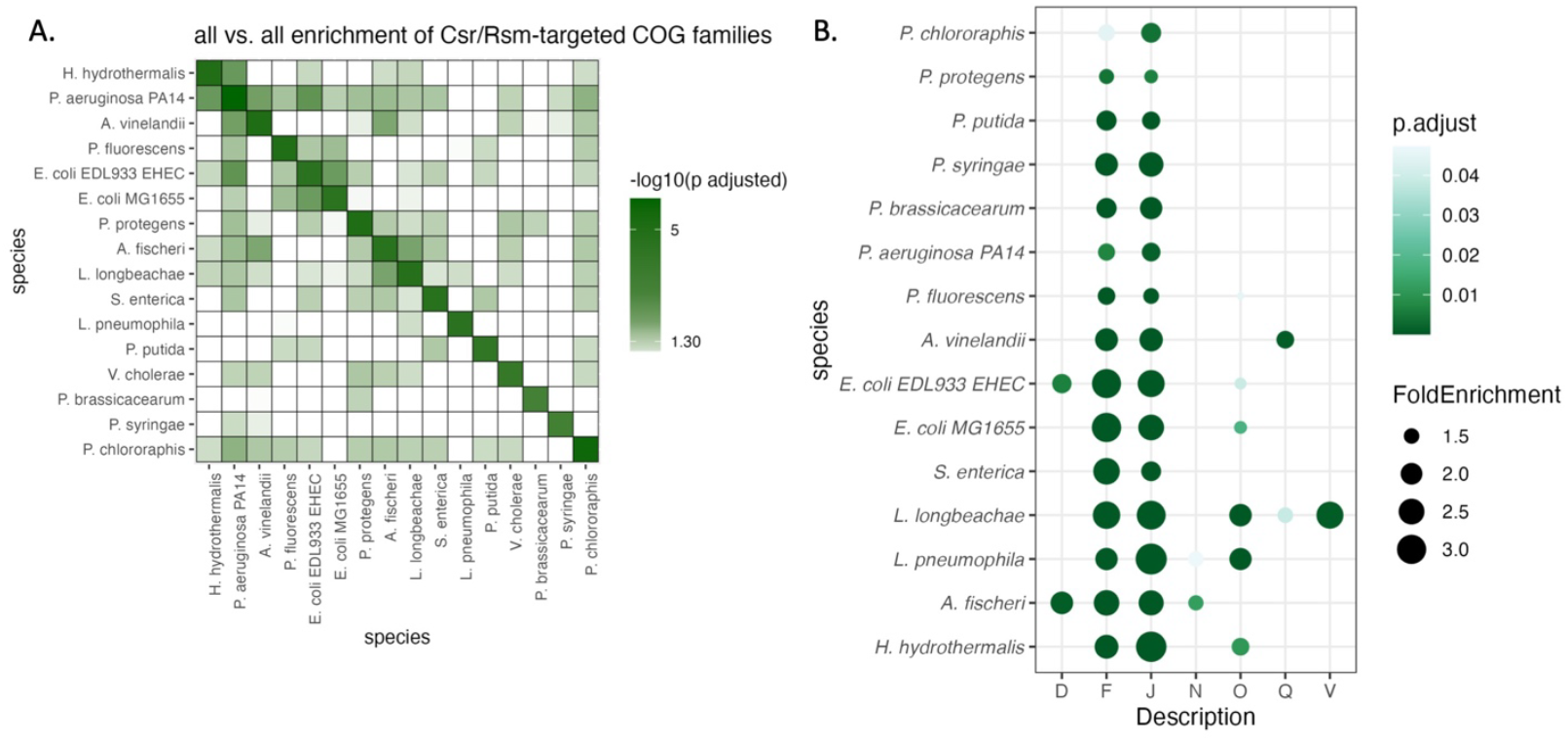
COG family enrichment varies across organisms modeled. A) All vs all overlap of enriched COG families shaded by significance of overlap, and clustered using hierarchical clustering. Not all predicted targets of CsrA converge onto the same families, and clustering does not follow the phylogenetic family level distribution observed on the orthogroup level. B) Enriched COG families per species modeled shows core overlap in processes such as F- nucleotide transport and metabolism. Family letter key is the following: D- Cell cycle control, division and chromosome partitioning, F- Nucleotide transport and metabolism, H- Coenzyme transport and metabolism, I – Lipid transport and metabolism, J- Translation, ribosomal structure and biogenesis, N- Cell motility, O- Posttranslational modification, protein turnover, and chaperones, Q- Secondary metabolite biosynthesis, transport and catabolism, V- Defense mechanisms.

GO enrichment analysis using TopGO [28]revealed a sharp contrast between pathogenic and nonpathogenic species (**Supplementary Figure 3**). Nonpathogens exhibited broader enrichment in membrane transport, stress response, and transposase activity, while pathogens were enriched in metal ion binding and protein-protein interaction functions. Only a few GO terms, such as those related to microcin transport and phosphonate metabolism, were shared across both groups.

Pfam domain analysis revealed that CsrA frequently targets transcriptional regulators, particularly response regulators (RRs) from two-component systems (**Supplementary Figure 4**). These targets were broadly conserved across species, suggesting that CsrA/RsmA exerts global regulatory influence by modulating key transcriptional hubs, including *qseB, arcA*, and *crp* in multiple organisms [7, 29, 30]. This regulatory architecture allows the CsrA/RsmA protein to control broad transcriptional programs without requiring direct binding sites in every mRNA present in each network.

Pathway enrichment analysis using the enrichKEGG() function from the ClusterProfiler package [31] revealed that while no single pathway was universally enriched, several trends emerged at the family level (**Figure 3**). Members of the Vibrionaceae and Enterobacteriaceae showed enrichment in secondary metabolite biosynthesis, while species-specific pathways included sulfur metabolism in *Aliivibrio fischeri*, xylene degradation in *Azotobacter vinelandii*, and chemotaxis in *Halomonas hydrothermalis*. GO enrichment analysis using TopGO [28]revealed a sharp contrast between pathogenic and nonpathogenic species (**Supplementary Figure 3**). Nonpathogens exhibited broader enrichment in membrane transport, stress response, and transposase activity, while pathogens were enriched in metal ion binding and protein-protein interaction functions. Only a few GO terms, such as those related to microcin transport and phosphonate metabolism, were shared across both groups. COG enrichment analysis showed that while some categories like nucleotide transport and metabolism (COG J, F) were commonly regulated, others were more niche-specific (**Figure 4A–B**). For example, *L. longbeachae* uniquely showed enrichment in defense mechanisms (COG V). Pfam domain analysis revealed that CsrA frequently targets transcriptional regulators, particularly response regulators (RRs) from two-component systems (**Supplementary Figure 3**). These targets were broadly conserved across species, suggesting that CsrA/RsmA exerts global regulatory influence by modulating key transcriptional hubs, including *qseB, arcA*, and *crp* in multiple organisms [7, 29, 30]. Together, these findings underscore the core and niche-specific reshaping of the CsrA/RsmA regulation: it maintains core functions across species while also adapting to specific environmental and physiological demands. This functional plasticity likely contributes to the ecological success and diversity of processes across γ-proteobacteria

### 2.5 Pathogenic and nonpathogenic E. coli strains show dramatic CsrA regulon divergence despite genome similarity

Given the observation that CsrA/RsmA regulation varies depending upon environmental niche, we examined intraspecific regulon divergence by comparing two E. coli strains that differ in pathogenicity. While nonpathogenic *E. coli K-12 MG1655* (EC MG1655) and enterohemorrhagic *E. coli O157:H7 EDL933* (EC EHEC) share a core set of 2,871 genes [32], only 270 were predicted to be overlapping CsrA targets, suggesting intraspecific divergence in post-transcriptional regulation (**Figure 5**).

**Figure 5:**
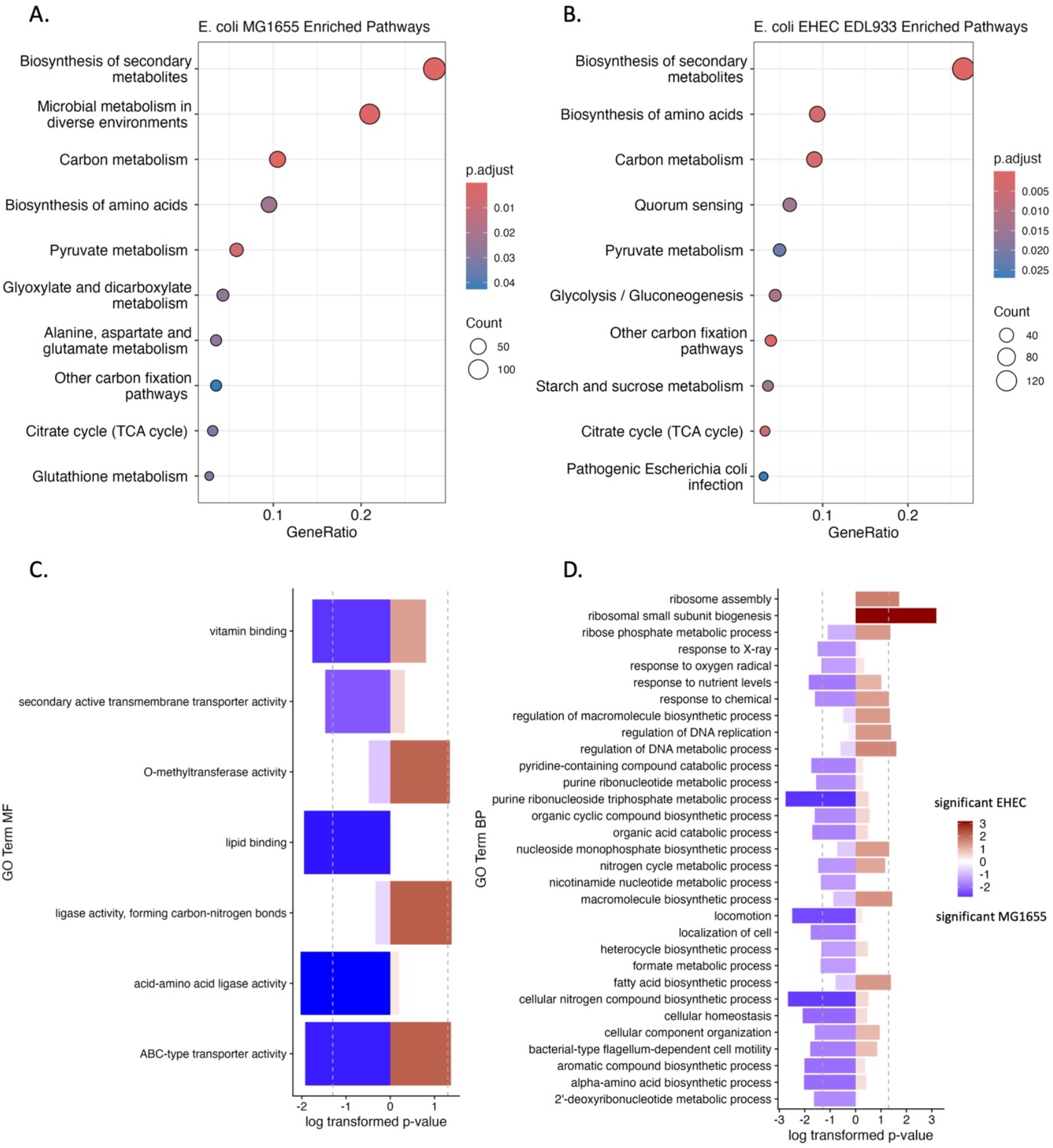
Shared and exclusive regulatory comparisons between predicted targets in *E. coli MG1655* and *E. coli O157:H7 EDL933 EHEC*. A-B) KEGG pathway enrichment for each variant modeled reveal shared and exclusive pathways enriched. C-D) GO term enrichment shows differentiation between regulated processes and functions.

In EC MG1655, CsrA targets were enriched in transmembrane transporter activity and general metabolic functions, consistent with regulation optimized for laboratory growth conditions. In contrast, EC EHEC showed enrichment in virulence-associated processes, including antibiotic resistance, biofilm formation, and quorum sensing. Notably, CsrA was predicted to regulate the Shiga toxin genes (*stxA, stxB*) and multiple LEE-encoded effectors (*tir, map, espF, espZ, cro, grlR, escE*) as well as non-LEE effectors (*nleA, nleB, nleD, nleF, nleH, espJ*) highlighting its role in pathogenicity [24, 33].

Both strains shared regulation of the response regulator *qseB*, part of the QseBC two-component system, which is involved in stress response and virulence [29]. This shared regulation underscores the dual role of CsrA in maintaining core regulatory functions while enabling strain-specific adaptations. The extent of regulon rewiring between these closely related strains demonstrates that post-transcriptional regulatory networks can evolve rapidly during niche adaptation.

### 2.6 Family-level analysis reveals lifestyle-specific CsrA/RsmA regulatory programs across γ-proteobacteria

Our observation of family-level enrichment of orthogroups (**Figure 3A**) prompted further detailed examination of regulatory patterns within phylogenetically related groups. We identified distinct regulatory patterns associated with specific lifestyles: plant-associated bacteria, pathogens with different host associations, and environmental specialists.

#### 2.6.1 Plant-associated Pseudomonads regulate nutrient cycling and biocontrol pathways

Within the highly diverse genus *Pseudomonas* [34, 35], plant-associated and soil-dwelling species show RsmA regulatory signatures of environmental adaptation. P. syringae and P. putida exhibit strong enrichment in glyoxylate and dicarboxylate metabolism, and metabolism in diverse environments (**Figure 3B**) consistent with their environmental versatility and roles in nutrient cycling [6, 36]. Secondary metabolite production and glutathione metabolism pathways were enriched in *P. putida* and *P. fluorescens*, species known for their biocontrol properties and interactions with plant hosts [37]. Notably, RsmA targets in *P. protegens* include genes involved in motility, secretion systems, and biofilms highlights adaptation to complex soil environments with diverse chemical stressors [38]. *A. vinelandii* shows unique enrichment in aromatic coumpound degradation: predicted CsrA targets include genes in xylene and benzoate degradation pathways relevant for bioremediation in diverse soil environments.

#### 2.6.2 Pathogenic P. aeruginosa shows enrichment in virulence and chronic infection pathways

In contrast to environmental pseudomonads, *P. aeruginosa* exhibits RsmA regulation of quorum sensing, biofilm formation, and phenazine biosynthesis (**Figure 3B**). These pathways are critical for establishing and maintaining chronic infections. This regulatory specialization reflects adaptation to the human host environment and aligns with known ΔrsmA mutant phenotypes showing altered biofilm formation and quorum sensing responses.

#### 2.6.3 Vibrionaceae species show host-specific regulatory adaptations

The CsrA regulons within the *Vibrioanaceae* exhibit unique host associated speciation. In *V. cholerae*, CsrA regulates a broad set of response regulators involved in virulence and environmental sensing, including *phoB, ompR, qseB, arcA, varA*, and *crp* [7, 30]. It also targets *toxR*, a master regulator of virulence, and *aspA*, involved in amino acid metabolism [39, 40]. These targets support the bacterium’s transition from aquatic environments to the human gut [41]. In *A. fischeri*, a symbiotic marine bacterium, CsrA regulates sulfur metabolism and luminescence. Predicted targets include *cysB, cysE, cysK, cysJ, cysI*, and *cobA*, which are essential for cysteine biosynthesis and symbiosis with the squid host *Euprymna scolopes* [42, 43]. These sulfur metabolism genes are not CsrA targets in *V. cholerae*, suggesting species-specific adaptation despite membership in the same family.

#### 2.6.4 Legionellaceae species regulate intracellular lifestyle genes with species-specific environmental adaptations

CsrA regulons in members of the *Legionellaceae* reflect their intracellular lifestyle while showing species-specific adaptations to different environments. In *Legionella pneumophila*, CsrA regulates numerous horizontally acquired genes of eukaryotic origin, including *ankH* and other ankyrin repeat proteins, many of which are associated with the Dot/Icm type IV secretion system [4, 44]. Predicted targets include *lpp06050, lpp2323, lpp2214*, and *lpp2038*, involved in poly-3-hydroxybutyrate (PHB) biosynthesis (**Supplementary Figure 5**), which supports metabolic shifts during intracellular replication [45, 46]. In *Legionella longbeachae*, CsrA targets differ, with regulation of *yneI* (aldehyde dehydrogenase) and *rlmI* (methyltransferase), suggesting adaptation to soil and compost environments [47]. Only three eukaryotic-origin genes: *icmE*, a calcium-transporting ATPase, and a cytokinin dehydrogenase are shared CsrA targets between the two species.

#### 2.6.5 The extremophile H. hydrothermalis shows CsrA regulation of specialized stress response pathways

The deep-sea extremophile *H. hydrothermalis* shows CsrA regulation of genes essential for survival in high-pressure, high-salinity environments, including chemotaxis (*MCP, Aer*), PHB biosynthesis (*phaA, phaB*), and starch/sucrose metabolism [48, 49]. CsrA regulation of *Aer* suggests a role for CsrA in oxygen sensing and metabolic switching in facultative anaerobic conditions [50]. Notably, the transcriptional activator phaR is not predicted to be bound by CsrA, suggesting CsrA controls PHB-mediated energy storage through post-transcriptional control rather than transcriptional mechanisms.

Together, these family-level analyses demonstrate that CsrA/RsmA regulation not only governs basic physiology but also fine-tunes traits critical for ecological fitness and host interactions. This likely reflects evolutionary rewiring driven by niche-specific pressures, horizontal gene transfer, and promoter evolution. CsrA/RsmA thus functions as a regulatory hub, integrating environmental signals into tailored gene expression programs that enhance survival and competitiveness in diverse habitats.

## 3.0 Discussion

Our comparative analysis across 16 γ-proteobacterial species reveals a fundamental principle of post-transcriptional regulatory evolution: conservation of molecular mechanism does not necessitate conservation of regulatory targets. Despite structural conservation of CsrA/RsmA (**Table 1**) and binding motif energetics (**Figure 1**), predicted regulons diverge dramatically across species. Only two orthogroups, those encoding for *rfbB* and *smpB*, are shared among nonpathogens, while pathogens have no exclusively regulated orthogroups (Section 2.3). This pattern demonstrates that post-transcriptional regulatory networks undergo species-specific rewiring to meet ecological demands, with the mechanisms and constraints differing fundamentally from those governing transcription factor evolution.

The short ANGGA recognition motif (5 nucleotides) embedded in hairpin structures [13, 14] creates an evolutionarily flexible landscape where UTR mutations can readily create or eliminate functional CsrA binding sites without compensatory changes in the protein. This differs from DNA-binding transcription factors, which require precise spacing relative to promoters and often co-evolve with their binding sites through compensatory mutations [51, 52]. CsrA’s positional flexibility across a target sequence facilitates rapid regulatory adaptation. A single UTR mutation forming an ANGGA-containing hairpin can place a gene under CsrA control without upstream regulatory architecture changes.

Our analysis of binding patterns in *A. vinelandii* supports this model: CsrA targets show significantly lower nucleotide diversity than non-targets (p = 0.014; **Supplementary Figure 6**), indicating purifying selection maintains functional binding sites once they provide fitness benefits. In other species, the absence of significant diversity differences suggests that while CsrA sites can emerge frequently by drift [53], their fixation depends on whether resulting regulation benefits that species’ niche. This evolutionary strategy where a broadly functional RNA-binding protein is conserved while rapidly evolving mRNAs contain its recognition motif may be more efficient for bacteria than protein-RNA co-evolution, particularly given short generation times, large population sizes, and strong selection pressures [54, 55].

Pathway and functional enrichment analyses reveal striking niche-specific patterns. Pathogens show enrichment in metal ion binding and protein-protein interaction functions, while nonpathogens exhibit broader regulation of membrane transport, stress response, and transposase activity (**Supplementary Figure 3**). The clearest example comes from *E. coli*: despite sharing 2,871 core genes [32] and recent divergence, MG1655 and EHEC O157:H7 strains show dramatically different CsrA regulons (**Figure 5**). In EHEC, our model predicts CsrA regulates Shiga toxin genes and Type III secretion system effectors, while MG1655 regulation focuses on transmembrane transport and general metabolism. This intraspecific divergence demonstrates that regulon rewiring occurs rapidly during niche adaptation. Acquisition of pathogenicity islands in EHEC was accompanied by integration of virulence factors into the existing CsrA network, suggesting active selection for coordinated virulence regulation. Family-level examples further illustrate niche adaptation. In *Legionella* species, CsrA regulates numerous horizontally acquired eukaryotic-origin genes [44]. *L. pneumophila* shows regulation of PHB biosynthesis genes supporting metabolic shifts during infection, while *L. longbeachae* regulates aldehyde dehydrogenase and methyltransferases relevant to environmental persistence. Despite close phylogenetic relationship and shared intracellular lifestyle, only three eukaryotic-origin genes are shared CsrA targets, demonstrating adaptation to subtle niche differences.

The family-level clustering of overlapping orthogroups (**Figure 3A**) indicates that while species-specific adaptation is extensive, some regulatory relationships persist at broader phylogenetic scales. This hierarchical pattern where core functions are conserved at family level but extensive divergence is preserved at the species level resembles a core-periphery regulatory architecture, where conserved central processes are surrounded by variable peripheral functions that adapt to specific ecological niches [56]. For CsrA/RsmA, the conserved function appears to be core metabolic processes (carbon metabolism, nucleotide metabolism exemplified by *relA* and *ftsZ*), while niche-specific processes (virulence, symbiosis, environmental stress) show extensive variation.

Other mechanisms likely drive regulon divergence, such as horizontal gene transfer and pathogenicity island integration. In EHEC, the LEE pathogenicity island contains genes with CsrA binding sites. Post-acquisition mutations likely created binding sites that were subsequently selected for coordinated virulence regulation, as LEE-like islands in other Enterobacteriaceae show different regulatory architectures. In addition, promoter and UTR evolution contributes to regulon divergence. The 100-200 bp 5’ UTRs have the evolutionary flexibility to mutate more rapidly than coding sequences due to weaker purifying selection. Mutations altering secondary structure can create or destroy CsrA binding sites. The *A. vinelandii* finding that targets show reduced diversity suggests purifying selection maintains sites once functionally important, implying that site creation occurs by drift but fixation depends on fitness benefits.

CsrA/RsmA likely expand their global regulatory control through the co-option of transcriptional regulators. CsrA frequently targets response regulators from two-component systems (**Supplementary Figure 4**), creating regulatory cascades where CsrA/RsmA indirectly influences entire transcriptional programs. The specific response regulators targeted differ by species (*qseB* in *E. coli* and *V. cholerae, toxR* in *V. cholerae*, multiple RRs in *P. aeruginosa*), suggesting repeated integration into existing transcriptional networks through evolution of binding sites in regulatory hubs. This “regulatory capture” allows CsrA to control niche-specific pathways efficiently without requiring binding sites in every structural gene.

CsrA regulation shows both similarities and differences compared to DNA-binding transcription factor evolution. Both show conserved binding mechanism but divergent target repertoires. However, CsrA binding sites face fewer positional constraints. Sites can be found anywhere in the 5’ UTR or early CDS with appropriate structure thus making them easier to evolve and potentially explaining more extensive regulon divergence compared to some TF families. The CsrA motif (5 bp ANGGA requiring structural context) differs fundamentally from typical TF sites (16-20 bp with moderate degeneracy). While the sequence motif is extremely common, functional sites are rarer due to structural requirements. This means sequence mutations alone can create or destroy potential sites, but actual regulatory impact depends on context. Additionally, CsrA can have different effects on different targets (stabilization vs. degradation, translational activation vs. repression) depending on binding site position relative to RBS and other structural features. This mechanistic flexibility may contribute to CsrA’s ability to regulate diverse processes, wherein the same protein can be repurposed for different regulatory outcomes simply by where binding sites evolve. Both TFs and CsrA show a pattern of “regulatory capture” of key hub genes (response regulators for CsrA, other TFs for transcription factors) to gain indirect pathway control. These parallels suggest some principles of regulatory network evolution may be universal across molecular mechanisms.

The patchy phylogenetic distribution of CsrA/RsmA across bacteria [34] suggests this regulatory system provides fitness benefits primarily under specific selective pressures. Our data suggest these include: frequent environmental transitions requiring rapid metabolic remodeling, pathogenic lifestyles requiring coordinate virulence regulation, and complex multi-stage life cycles (free-living vs. host-associated states). As noted in the introduction, CsrA/RsmA exhibits “lifestyle-adaptive distribution” it confers fitness advantages specifically in organisms requiring rapid, coordinated metabolic or behavioral switches. In more stable environments or in organisms with specialized metabolisms, alternative regulatory strategies may suffice.

Several important questions emerge from this work: (1) Systematic CLIP-seq across the modeled species would provide ground truth for predictions and reveal species-specific binding features. (2) Future models should integrate sponge sRNA effects on CsrA/RsmA activity. (3) Population genomic approaches comparing closely related strains could reveal binding site gain/loss rates and signatures of selection. (4) Laboratory evolution experiments in different environments with and without functional CsrA/RsmA could test whether regulon rewiring occurs on observable timescales. (5) Extending this approach to α-proteobacteria, other γ-proteobacterial families, and Gram-positive bacteria that encode for CsrA/RsmA family proteins would reveal whether observed patterns hold more generally.

## 4.0 Conclusions

In this work, we detail the process of evaluating, validating, and predicting Csr/Rsm binding to a selection of 16 species across the γ-proteobacteria. These species represented organisms with varying degree of pathogenicity and are found in different environments. In applying the model to each organism, we identify that CsrA/RsmA binding mechanism is conserved across these species, however, the regulatory program is not.

This work demonstrates that the model is generalizable to organisms beyond *E. coli* and *P. aeruginosa*. This indicates that the underlying principles of CsrA/RsmA binding is applicable to a broader range of γ-proteobacteria, making this process a useful tool in studying impact of the global regulator in less-characterized organisms. Modeling is useful for generating new hypotheses regarding the regulatory network of CsrA in other organisms. Each species had predictions of CsrA targeting exclusive and overlapping pathways, highlighting how post-transcriptional regulation changes to adapt to different environments.

## 5.0 Materials and Methods

### 5.1 Selection methods

CsrA protein sequences were extracted from each organism and PDB structures of the CsrA monomeric sequence in complex with the RsmZ hairpin loop “CCCCGAAGGAUCGGGG” were generated using AlphaFold 3 [21]. PDB files were generated from these predicted structures using the PyMol Molecular Graphics System verion 3.0.3. Position Weight Matrices were generated per species using these PDB structures as input into the RNP ΔΔG tool on the Rosie webserver using default parameters [23] and analyzed as described in section 2.1.

UTR sequences modeled were extracted from all 16 reference genomes (**Supplementary Table 4**) based on TSS mapping, where available. This ensures that our model accurately reflects the transcriptional landscapes of the studied organisms, allowing for more precise predictions of CsrA binding and regulatory effects. Where TSS mapping was not available, 100 bases preceding and following the start codon were selected to be used in the model. These sequences were sliced from the corresponding genome fasta file using custom scripts written in Python 3.9, or in R 4.4.1. Modeling performed on the Stampede2 compute cluster within the Texas Advanced Computing Center.

### 5.2 Position Weight Matrix Generation and Modeling

Given the structural similarity and high correlation between PWM affinities for specific motifs, we applied the same biophysical modeling scheme as described in [12]. The 5’ UTR and first 100 bases of coding sequence were evaluated using the same model and position weight matrix crafted for RsmA. Transcriptomes were extracted from reference genomes for each species in **Table 2**.

**Table 2.**
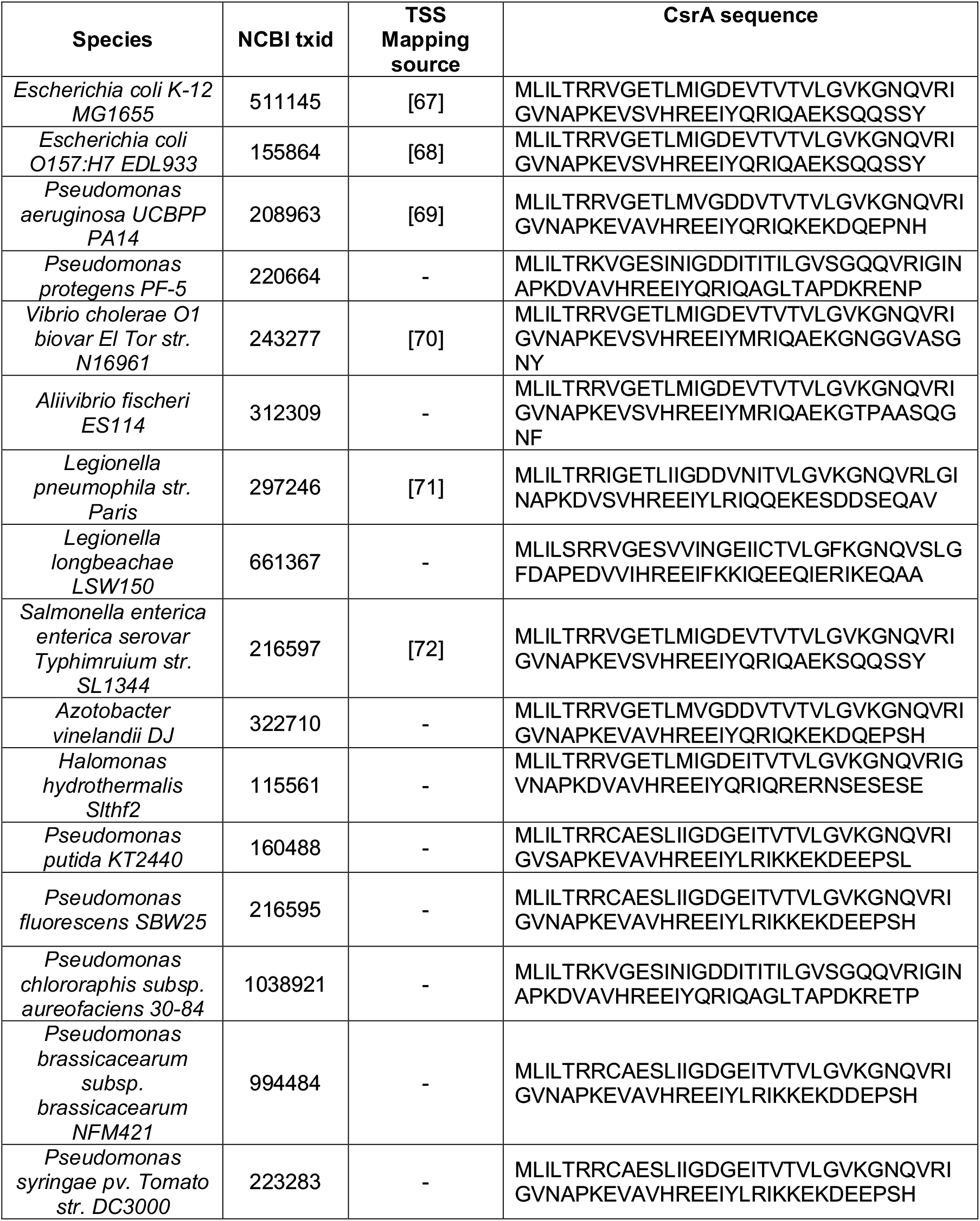
Summary of species modeled in this work. If TSS were mapped, the source is provided. Leader sequences from species with no TSS sources were extracted -100:+100 surrounding the start codon as defined in the feature table downloaded from NCBI.

### 5.3 Scoring Diversity

Nucleotide diversity for all leader sequences modeled were calculated using the Pairwise Alignment Positional Nucleotide Counting (PAPNC) method [57]. Modeled sequences were grouped by species and prediction. Frequency of each nucleotide were calculated using custom scripts in Python 3.11.5 and visualized in R 4.4.1. To test for significant differences in nucleotide diversity, a generalized additive model (GAM) was fit to our datasets using the R package, mgcv 1.9-1. In this model, diversity was selected as a response variable, the position as the smoothing term, and model prediction (target/non-target) as an additional predictor variable.

### 5.4 Orthogroup mapping and statistical analysis

To map protein coding genes to orthogroups, the EggNOG-mapper v2 tool was used on each genome protein coding data using the default parameters. Aggregation of this annotation data was aggregated in R. The enricher() function within the ClusterProfiler package was used to test for specific enrichment of KEGG orthology, PFAM domains, and COG families within individual species. Enricher() was used with the following parameters: p-adjust < 0.05, Benjamini-Hochberg error correction.

Overlap enrichment analyses for overlapping orthogroups, COG families, and PFAM clans were performed using custom scripts that utilize the phyper() function in R. Pathway enrichment analyses were performed in R using ClusterProfiler v4.12.6 and topGO v2.56.0. All additional statistical analysis of model data was performed in R 4.4.1.

## Acknowledgements

We thank Dr. Phillip Sweet for his advice and considerations in cross-species analysis. We also thank Dr. Matthew Burroughs, Kobe Grismore, Ryan Buchser, and Agata Turula for their feedback during the preparation of this manuscript.

## Author Contributions

AJL: Conceptualization, Data curation, Formal Analysis, Funding acquisition, Investigation, Methodology, Software, Validation, Visualization, Writing–original draft, Writing–review and editing. LGH: Data curation, Analysis all L. pneumophila transcriptome data. AS: Prepared and analyzed the S. enterica data. LMC: Conceptualization, Funding acquisition, Supervision, Writing–original draft, Writing–review and editing.

## Data access

Scripts for model and associated files can be found at https://github.com/ajlukasiewicz/rsm_biophysical_model.

## Ethics Statement

This computational study utilized only publicly available genomic sequences and previously published experimental datasets. No human subjects, animal subjects, or field sampling was involved in this research. Therefore, no institutional review board approval or ethics committee oversight was required.

