## Supplementary Figures for "Modeling binding of the conserved Csr/Rsm protein family across species of the γ-proteobacteria reveals niche-specific adaptation of the post-transcriptional regulon"

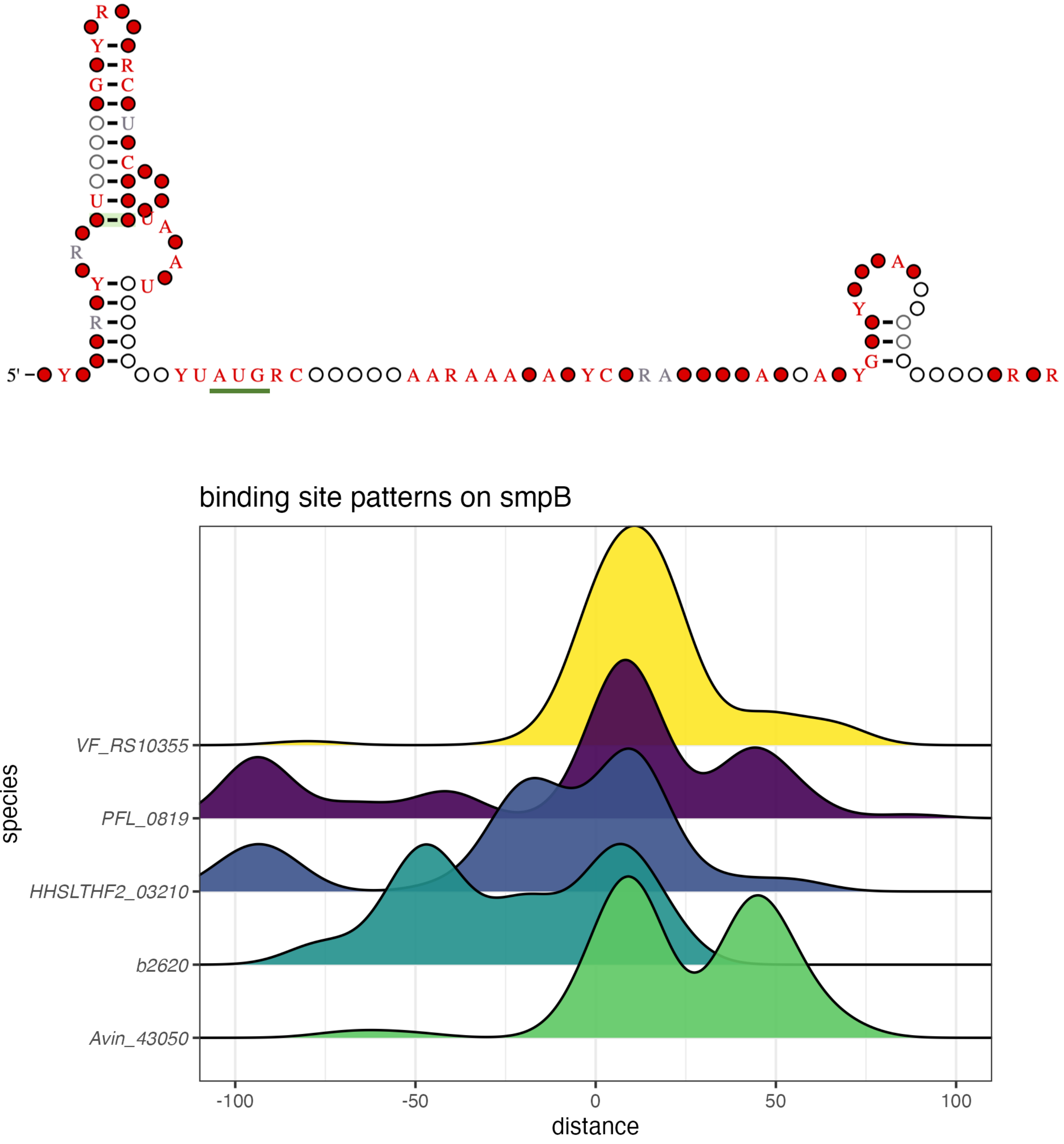


**Supplementary Figure 1: structure and predicted binding sites for nonpathogens encoding *smpB* 5' UTR.** A) R-scape generated secondary structure conservation for the *smpB* leader sequence modeled. The start codon is underlined in green. B) Binding site peak distributions for the leader sequence in each nonpathogen modeled. Interestingly, the highest frequency of site predictions fall at the start codon and extend into the CDS, which falls along the position of the two conserved structured elements within *smpB.*

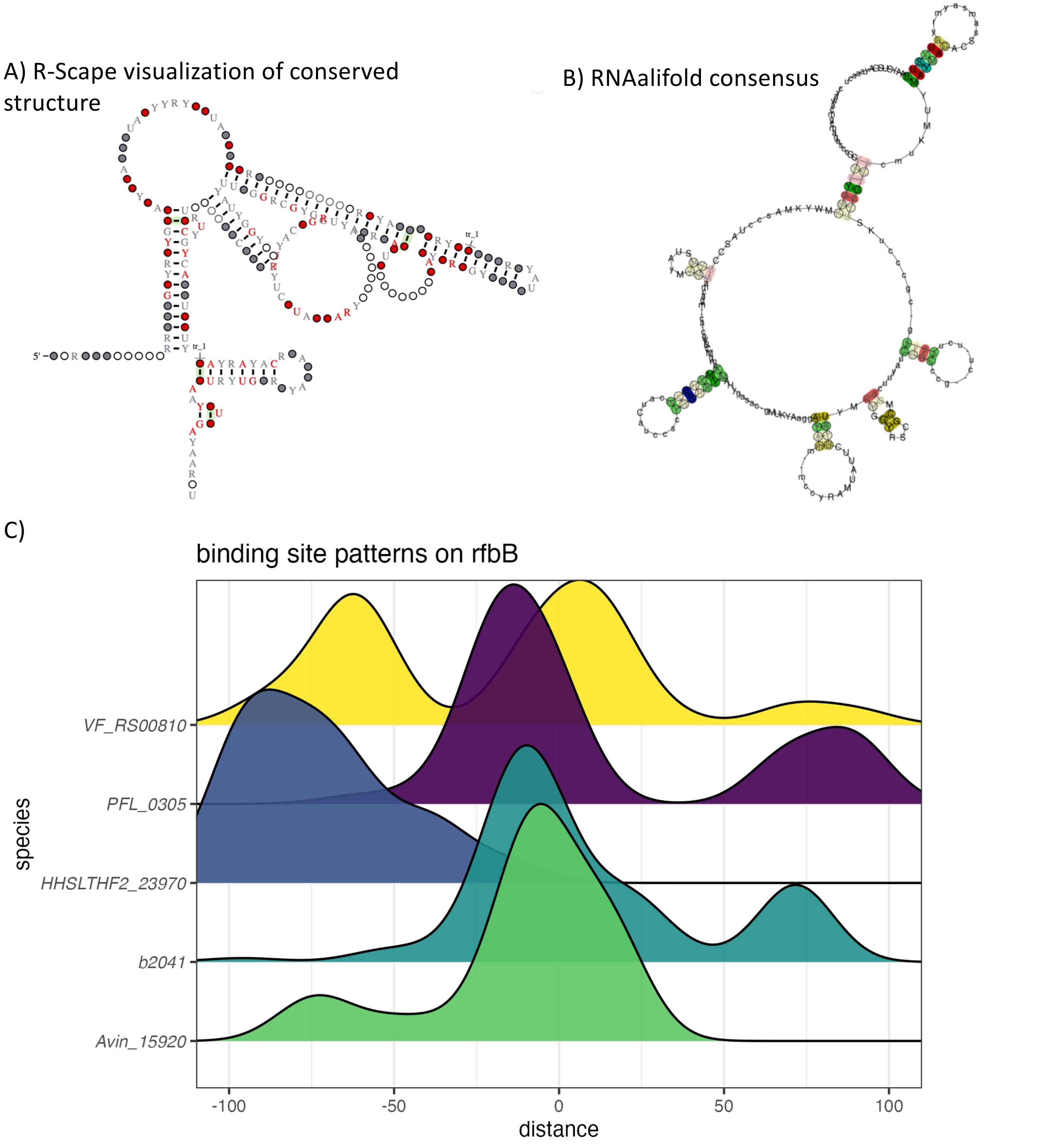


**Supplementary Figure 2: structure and predicted binding sites for nonpathogens encoding rfbB 5' UTR.** A) R-scape generated secondary structure conservation for the *rfbB* leader sequence modeled. B) RNAalifold consensus structure of the leader sequence shows cconserved stem loops surrounding the start codon as well. C) Binding site peak distributions for the leader sequence in each nonpathogen modeled. Interestingly, binding peak distributions in *H. hydrothermalis* fall further upstream of the start codon, as observed for the other nonpathogens. .

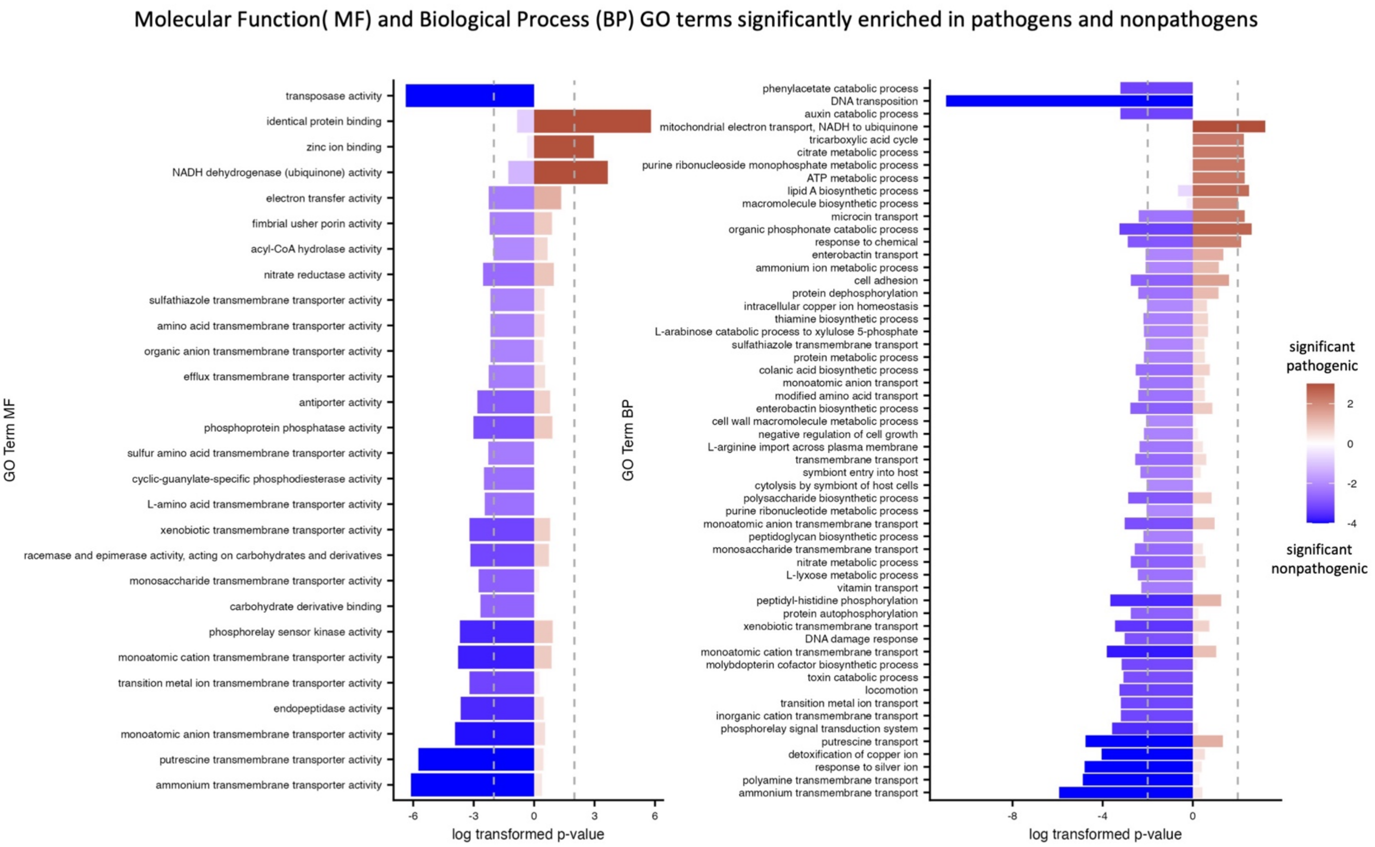


**Supplementary Figure 3: Significantly enriched Molecular Function (MF; left) and Biological Process (BP; right )**

GO terms differ between pathogenic (red) and nonpathogenic (blue) bacterial types modeled. A minority of GO terms are shared between the two types, and CsrA appears to regulate processes more broadly in nonpathogens relative to pathogens.

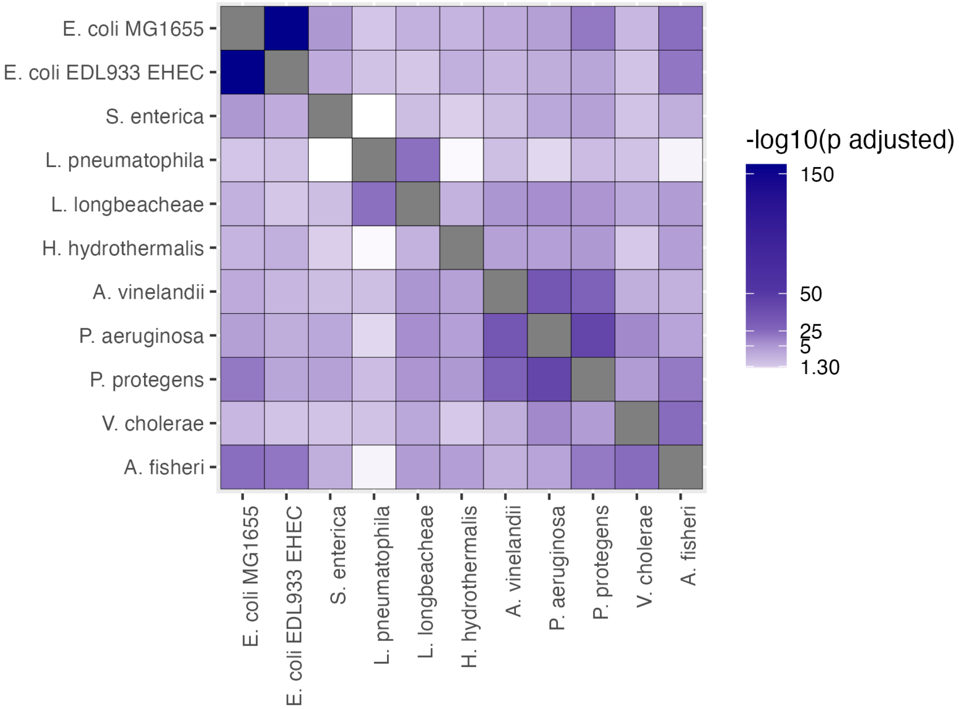


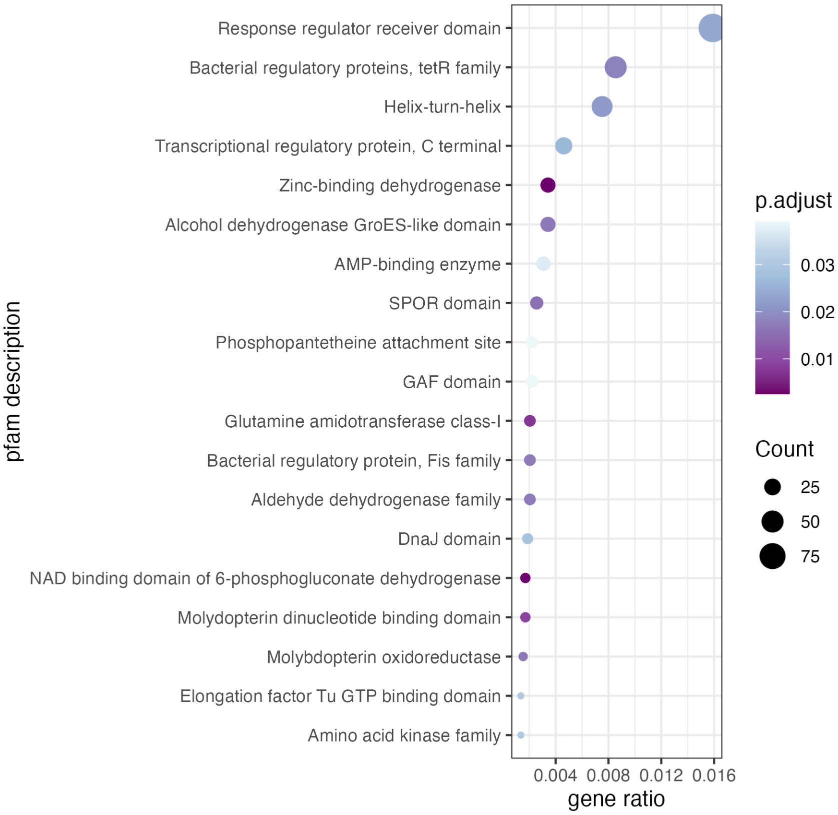


**Supplementary Figure 4: A**) All vs all overlap of enriched PFAM domains shaded by significance of overlap and clustered using hierarchical clustering. B) Targeted domains converge onto key transcriptional regulatory nodes that likely influence global transcriptional shifts in the organism.

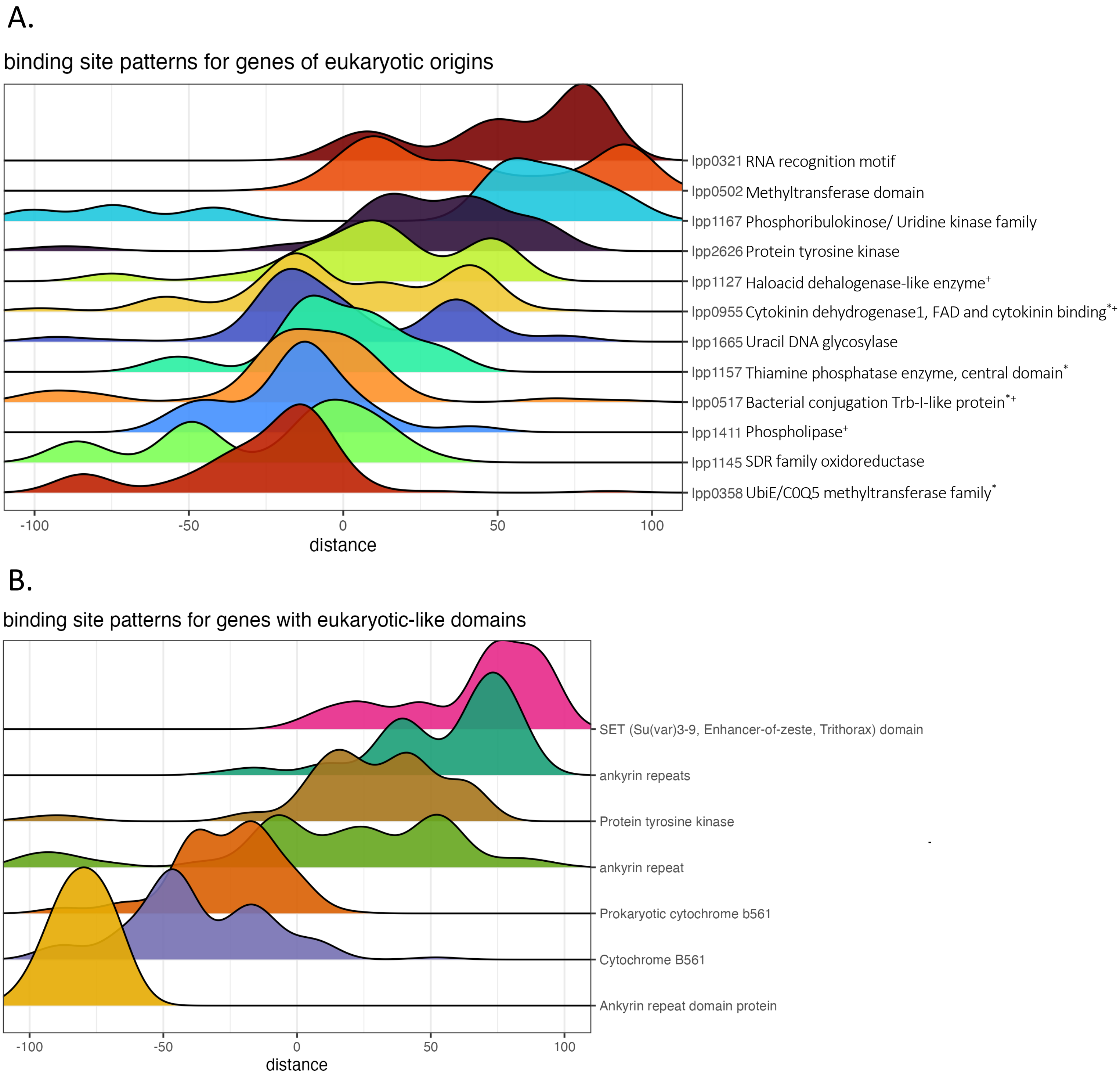


**Supplementary Figure 5: Binding site patterns for genes of eukaryotic origins and eukaryotic-like domains predicted to be bound by CsrA in *L. pneumophila*.** A) Distribution of predicted binding sites across genes of eukaryotic origin. Asterisk indicates genes with prior evidence of CsrA binding (Sahr et. al., 2018), Cross indicates gene is also predicted to be bound by CsrA in *L. longbeachae.* B) Binding site patterns for genes with eukaryotic-like domains that span far upstream of the 5' UTR and into the CDS. These include ankyrin repeat proteins such as *ankH.*

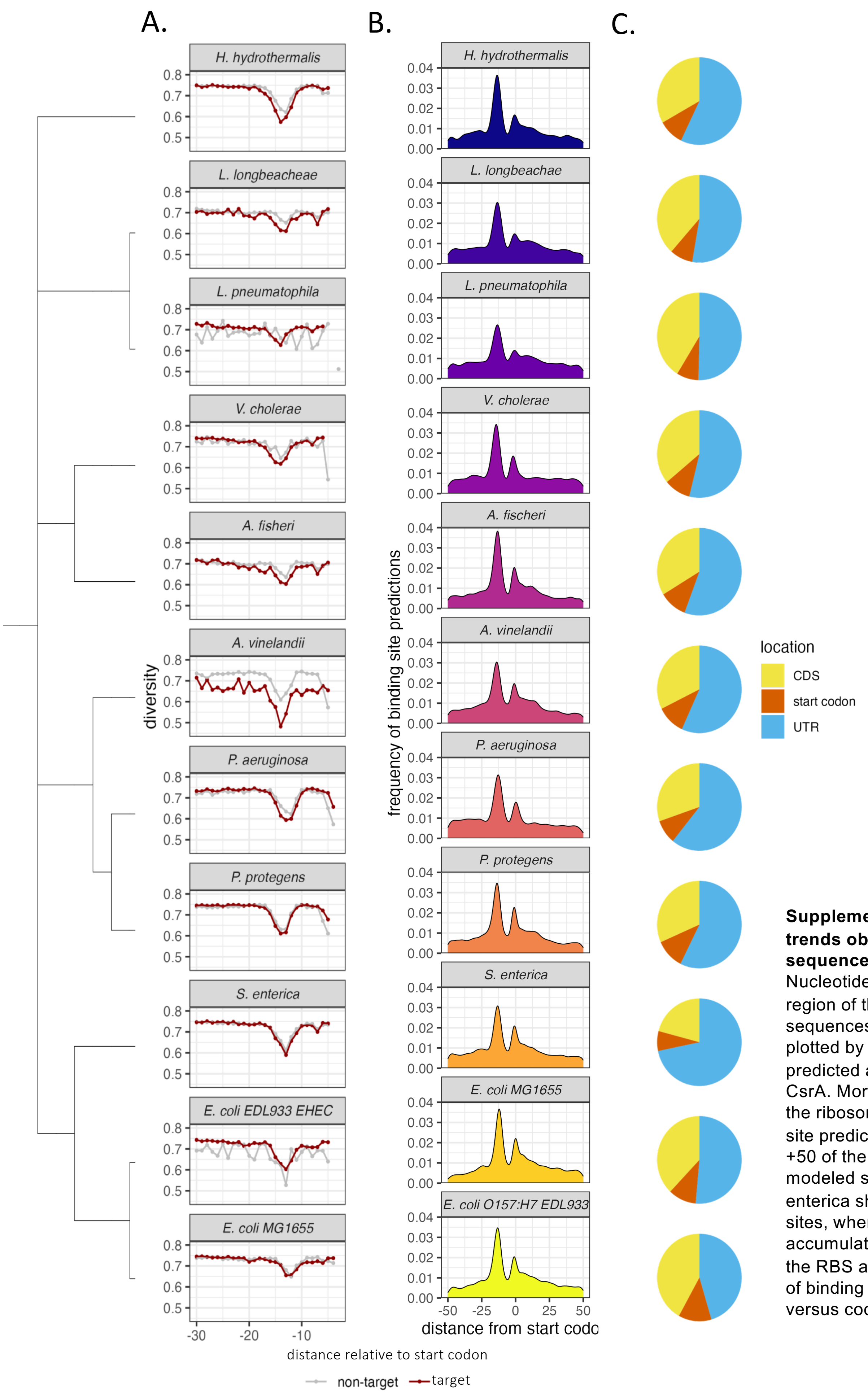


**Supplementary Figure 6: Molecular trends observed for modeled UTR sequences across 11 species.** A) Nucleotide diversity in the -30 – 0 region of the 5' UTRs of all modeled sequences. Nucleotide diversity is plotted by whether the genes were predicted as targets or non-targets of CsrA. More conservation surrounding the ribosome binding site. B) Binding site prediction frequencies from -50 to +50 of the start codon across the 6 modeled species. Predictions for S. enterica shows a broader distribution of sites, whereas the other species accumulate predicted binding peaks at the RBS and start codon. C) Proportion of binding sites predicted in UTR versus coding region.
